# Assessment of non-linear mixed effects model-based approaches to test for drug effect using simulated data: type I error and power properties

**DOI:** 10.1101/2024.04.13.589388

**Authors:** E. Chasseloup, A. Tessier, M.O. Karlsson

## Abstract

Pharmacometric approaches achieves higher power to detect a drug effect compared to traditional statistical hypothesis tests. Known drawbacks come from the model building process where multiple testing and model misspecification are major causes for type I error inflation. IMA is a new approach using mixture models and the likelihood ratio test (LRT) to test for drug effect. It previously showed type I error control and unbiased drug estimates in the context of two-arms balanced designs using real placebo data, in comparison to the standard approach (STD). The aim of this study was to extend the assessment of IMA and STD regarding type I error, power, and bias in the drug effect estimates under various types of model misspecification, with or without LRT calibration. Two classical statistical approaches, t-test and Mixed-Effect Model Repeated Measure (MMRM), were also added to the comparison. The focus was a simulation study where the extent of the model misspecification is known, using a response model with or without drug effect as motivating example in two sample size scenarios.

The IMA performances were overall not impacted by the sample size or the LRT calibration, contrary to STD which had better type I error results with the larger sample size and calibrated LRT. In terms of power STD required LRT calibration to outperform IMA. T-test and MMRM had both controlled type I error. The t-test had a lower power than both STD and IMA while MMRM had power predictions similar to IMA. IMA and STD had similarly unbiased drug effect estimates, with few exceptions.

IMA showed again encouraging performances (type I error control and unbiased drug estimates) and presented reasonable power predictions. The IMA performances were overall more robust towards model mis-specification compared to STD. IMA confirmed its status of promising NLMEM-based approach for hypothesis testing of the drug effect and could be used in the future, after further evaluations, as primary analysis in confirmatory trials.

## 2 Introduction

The increase in power to detect a drug effect with pharmacometric model-based approaches and mixed-effect models compared to conventional statistical hypothesis tests was largely documented[1–7]. While there are consequently widely used in exploratory trials with learning purposes, their use remains limited in confirmatory trials[8]. In a review of 198 submissions to the Food and Drug Administration over the years 2000 to 2008 [8], pharmacometric analyses contributed to drug approval and labeling decisions for more than 60% of submissions, but only five (2.5%) used model-based primary endpoints in confirmatory trials, all concerning pediatric development. Type I error inflation and model misspecification are critical hurdle for theses approaches, and the risk of false positive is of major concern for decision making, as it can result in the pursuit of the development of an unsuccessful drug candidate, or the approval of an inefficient drug by the regulatory agencies. The statistics used to analyse such trials are hence critical.

To promote the use of pharmacometric models as primary analysis in confirmatory trials such as proof-of-concept studies, it is required to develop approaches that could result in a better type I error control. The use of pre-specified models through model-averaging method has been proposed and seems successful to control type I error rate, but there is little experience on its use in practice. Instead of the model building process resulting in the selection of a single model, model-averaging weights the model predicted quantity of interest based on a goodness-of-fit measure of a pre-selected set of models[9–11]. A promising alternative is the use of IMA, a novel pharmacometrics model-based approach using a mixture model[12], and using the LRT to test for drug effect. In IMA the population is assumed to be partitioned into a set of mixtures, i.e. sub-populations, described by different model features[13]. Mixture models are usually used to describe multimodal parameter distributions when no predictor explaining the observed data heterogeneity is available.

The present work aimed at extending the assessment of IMA to a simulation study where the extent of the model misspecification is controlled. The assessment was carried out with respect to type I error, power, and bias in the drug estimates, when testing for drug effect, under a small sample size and a large sample size scenario. STD and two traditional statistical approaches, T-test and MMRM (a longitudinal likelihood-based approach), were also assessed in the same framework. A drug-response model with or without drug effect was used as a motivating example to address clinical trial simulations.

## 3 Methods

NONMEM version 7.4.3[14] and PsN version 4.8.8[15] automated the simulation of clinical trials with or without drug effect, and the subsequent model estimation through the Stochastic Simulation and Estimation (SSE) functionality of PsN. The FOCE estimation method without interaction was used as the residual error model was additive. The PsN functionality for randomization tests (randtest) automated the calibration of the Likelihood Ratio Test (LRT). The processing of the results and the traditional statistical hypothesis tests were performed in the statistical software R version 3.5.2[16] and the lme4 package[17].

### 3.1 Study design

Clinical trials were simulated under a balanced (1:1 randomization) two-arms parallel design, with subjects receiving either a single dose of an active treatment or a placebo control. Each subject had four observations at t=0 (pre-dose), 1, 2, and 3 time units after the treatment initiation. A typical POC design scenario was simulated with 30 subjects per arm (60 in total), and a BIG design scenario with 120 subjects per arm (240 in total) to get conditions closer to the asymptotic properties of the LRT.

### 3.2 Simulation model

The data simulated corresponded to a longitudinal response composed of four elements: a baseline value, a placebo effect, a drug effect, and a residual error term (Eqn. 1).

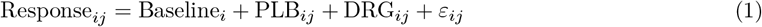

For individual *i* at time *j*, Response_*ij*_ is the observed response, Baseline_*i*_ is the response value observed at pre-dose, PLB_*ij*_ is the placebo effect, and DRG_*ij*_ is the corresponding drug effect component (with DRG_*i*0_=0). *ε*_*ij*_ ∼ 𝒩 (0, *σ*^2^) is an additive residual error term, *σ*^2^ being the variance. PLB_*ij*_ was described using a time-exponential placebo model (Eqn. 2), and DRG_*ij*_ was described using an offset model (Eqn. 3).

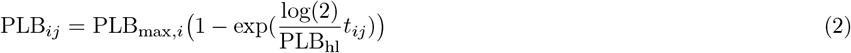

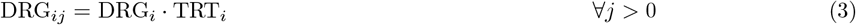

Where *t*_*ij*_ is the time of the observation *j* for individual *i*, PLB_max,*i*_ is the individual maximal placebo effect, PLB_hl_ is the half-life to achieve this maximal effect, DRG_*i*_ is the individual drug effect, and *TRT*_*i*_ is the variable describing the individual treatment allocation, equal to 0 for the placebo arm, and equal to 1 for the treated arm.

As indicated by the _*i*_ index in the equations above, the following parameters had between-subject variability (BSV): Baseline_*i*_, PLB_max,*i*_, and DRG modeled as random variables *η* ∼ 𝒩 (0, Ω). Ω being a variance-covariance matrix of the random effects where the diagonal elements 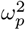 correspond to the variance of the *p*^*th*^ parameter. The individual parameters were log-normally distributed for Baseline and DRG (Eqn. 4), and normally distributed for PLB_max_ with random effect included in a multiplicative manner allowing negative values (Eqn. 5).

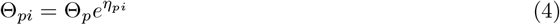

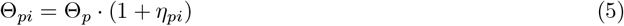

Where Θ_*pi*_ represents the value of the *p*_*th*_ parameter for individual *i*, Θ_*p*_ is the typical value and *η*_*pi*_ the value of the random effect of the *p*_*th*_ parameter for individual *i*. A covariance was introduced between *η*_Baseline_ and 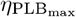. The values used in the simulations for the structural and statistical parameters presented above are provided in Table 2.

**Table 1:** Values of the parameters used for the clinical trial simulations.

| Parameters | Description | Value |
| --- | --- | --- |
| $\Theta_{\text{Baseline}}$ | Typical response value at pre-dose | 100 |
| $\Theta_{\text{PLB}_{\text{max}}}$ | Typical maximal placebo effect | 25 |
| $\Theta_{\text{PLB}_{\text{hl}}}$ | Typical half-life of the maximal placebo effect | 0.75 |
| $\Theta_{\text{DRG}}$ | Typical drug effect | 9 |
| $\omega_{\text{Baseline}}^2$ | Baseline<sub>*i*</sub> variance | 0.25 |
| $\omega_{\text{PLB}_{\text{max}}}^2$ | PLB<sub>max,*i*</sub> variance | 0.25 |
| $\text{cov}(\eta_{\text{Baseline}_i}, \eta_{\text{PLB}_{\text{max},i}})$ | Baseline<sub>*i*</sub>-PLB<sub>max,*i*</sub> covariance | -0.1 |
| $\omega_{\text{DRG}}^2$ | DRG<sub>*i*</sub> variance | 0.25 |
| $\sigma^2$ | Residual error variance | 100 |

**Table 2:** Name and description of response models fitted to simulated data.

| Model | Misspecification | Model features | Equation |
| --- | --- | --- | --- |
| *M1* | No misspecification | Delayed asymptotic placebo model, offset drug model | Identical to the simulation model |
| *M2* | Structural placebo model reduction | Offset placebo model | $PLB_{ij} = PLB_{\max,i}$ |
| *M3* | Structural placebo model reduction | Time-linear placebo model | $PLB_{ij} = PLB_{\text{slope},i} t_{ij}$ |
| *M4* | Structural placebo model extension | Weibull placebo model | $PLB_{ij} = PLB_{\max,i} (1 - \exp(\frac{\log(2)}{PLB_{hl}} t_{ij})^{PLB_{\text{power}}})$ |
| *M5* | Different structural placebo model | Power placebo model | $PLB_{ij} = PLB_{\text{slope},i} t_{ij}^{PLB_{\text{power}}}$ |
| *M6* | Placebo BSV model reduction | No covariance between Baseline and $PLB_{\max}$ | $\text{Cov}(\eta_{\text{Baseline}}, \eta_{PLB_{\max}}) = 0$ |
| *M7* | Placebo BSV model reduction | No BSV on $PLB_{\max}$ | $PLB_{ij} = PLB_{\max} (1 - \exp(\frac{\log(2)}{PLB_{hl}} t_{ij}))$ |
| *M8* | Placebo BSV model extension | Addition of BSV on $PLB_{hl}$ | $PLB_{ij} = PLB_{\max,i} (1 - \exp(\frac{\log(2)}{PLB_{hl,i}} \cdot t_{ij}))$ |
| *M9* | Different BSV placebo model | Log-normal distribution for $PLB_{\max,i}$ | $PLB_{\max,i} = PLB_{\max} e^{\eta_i}$ |
| *M10* | Different BSV placebo model | BSV on $PLB_{hl}$ instead of $PLB_{\max}$ | $PLB_{ij} = PLB_{\max} (1 - \exp(\frac{\log(2)}{PLB_{hl,i}} t_{ij}))$ |
| *M11* | Different structural drug model | Time-linear drug model | $DRG_{ij} = DRG_{\text{slope},i} t_{ij}$ |
| *M12* | Drug BSV model extension | Covariance between Baseline and DRG | $\text{Cov}(\eta_{\text{Baseline}}, \eta_{\text{DRG}})$ |
| *M13* | Different structural drug model | Delayed asymptotic drug model | $DRG_{ij} = DRG_{\max,i} (1 - \exp(\frac{\log(2)}{DRG_{hl}} t_{ij}))$ |

BSV: between-subject variability.

### 3.3 Estimation models

Thirteen different response models were fitted to the simulated data corresponding to different degrees of misspecification compared to the simulation model: reductions, extensions, or changes in the model structure or in the statistical model. The models are summarized in Table 2. The model *M* 1 had no misspecification, the models *M*2 to *M*5 introduced misspecifications in the structural placebo model, the models *M*6 to *M*10 introduced misspecifications in the BSV placebo model, and the models *M*11 to *M*13 introduced misspecifications in the drug model. The model *M*1 was identical to the model used for the simulations, hence assumed to be the “true” model. In *M*2 and *M*3 the placebo model was reduced respectively to an offset or a time-linear model. In *M*4 the placebo model was extended to a Weibull model while in *M*5 a different structure was used: a power model. The *M*6 and *M*7 corresponded both to BSV placebo model reductions by omitting respectively the covariance between *η*_Baseline_ and 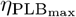, or the BSV of PLB_max_. *M*8 extended the placebo BSV model by adding a BSV distribution to the PLB_hl_ parameter and *M*9 assumed an exponential model for the BSV of PLB_max_ instead of the multiplicative one. *M*10 assumed BSV on PLB_hl_ instead of PLB_max_. *M*12 extended the BSV drug model by adding a covariance term between *η*_Baseline_ and *η*_DRG_. Both *M*11 and *M*11 assumed different drug model structure, respectively time-linear or delayed asymptotic.

### 3.4 Description of the approaches

Each of the thirteen models were declined in a null hypothesis (H0) and an alternative hypothesis (H1), differing on whether the treatment information was used in order to test for its significance to improve the data description using the LRT. The LRT compares two nested models to draw statistical conclusion on whether the feature added in the bigger model improves the description of the data. It uses the difference in OFV (Δ*OFV*) between the two models as test statistic, which is assumed to follow a *χ*^2^ distribution of *n* degrees of freedom when the models are nested, *n* being the number of additional parameters estimated in the alternative hypothesis. The hypotheses were different between STD and IMA.

In STD H0 the response model was fitted without drug effect, the drug effect parameters being estimated on the individuals allocated to the treated arm (TRT=1) only in H1. The number of degrees of freedom (df) for the LRT was set to the number of parameters of the drug model, corresponding to the additional parameters estimated in H1 (i.e. 2 df for the models *M*1 to *M*11 and 3 df for the models *M*12 and *M*13).

IMA assumed that the data was composed of two sub-populations (mixtures), the treated individuals and the non-treated individuals. Each of the sub-population was then described using a different submodel: a response model without drug effect or a response model with drug effect, corresponding respectively to Eqn. 1 with DRG_*ij*_ term or without. Accordingly the two submodels were fitted to the data both in H0 and in H1 using a mixture model. The two hypotheses differed by the estimation of the mixture proportion which was fixed to the placebo allocation ratio in H0, and estimated based on the treatment allocation in H1. Thanks to the mixture model, each submodel is fitted to each individual and their respective contribution to the *OFV*_*i*_ is estimated according to[18]:

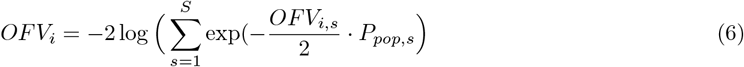

Where *OFV*_*i,s*_ is the individual objective function value for the sub-population *s* (*s* = 1, 2 in the present analysis), and *P*_*pop,s*_ is the mixture probability for the sub-population *s*. In the case of a balanced two-arm design in H0 every individual could be allocated to each of the submodel with the same probability (*P*_*pop*,1_=*P*_*pop*,2_=0.5). In H1, when the sub-population 1 corresponded to the treated individuals, the population probability *P*_*pop*,1_ was estimated according to Eqn. 7:

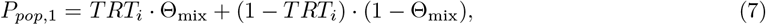

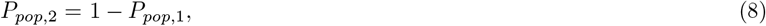

where Θ_mix_ is the mixture proportion, and *TRT*_*i*_ is the treatment allocation information. With IMA the number of df for the LRT was equal to 1 regardless of the drug effect model, as Θ_mix_ is the only additional parameter estimated in H1 compared to H0. When Θ_mix_ = 1 the *OFV*_*i*_ of the treated patients had no contribution from the submodel without drug effect. In that case IMA is equivalent to the STD approach.

Traditional statistical analysis, a t-test and MMRM, were also performed. The t-test tested for a significant difference between the two arms in the change from baseline at the last observation (*t* = 3). MMRM used two nested linear mixed effects models to describe the longitudinal individual response change from baseline CFB_*ij*_ according to Eqn. 9:

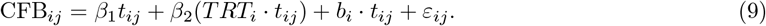

Here *β*_1_ is the time fixed effect and *β*_2_ quantifies the interaction between the treatment allocation and time. *β*_2_ is introduced in H1 only to test the assumption of a different average slope between the treatment groups. *b*_*i*_ is the random effects on the slope and *ε*_*ij*_ the residual error, both normally distributed around 0. No intercept was estimated as CFB_*i*0_ = 0 when *t* = 0.

### 3.5 Simulated data and evaluation

*N* = 500 datasets were simulated for the POC and the BIG design scenarios, both under H0 and H1 to mimick *N* trials without drug effect or with drug effect respectively. To quantify in the simulations the difference in response introduced by the drug effect, the change from baseline (CFB) was computed for each patient *i* in each dataset (*n* = 1, …, *N*), as the difference between each observation and its corresponding pre-dose observation. The placebo-corrected change from baseline of the response under treatment (ΔΔResponse) was then computed according to Eqn. 10:

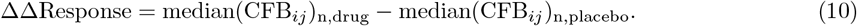

The type I error rate (concluding falsely to the presence of a drug effect) and the power predictions (rejecting falsely the presence of a drug effect) were computed for each approach, each scenario, and each of the thirteen models presented above. The type I error rate was defined as the fraction of the *N* datasets for which H0 was rejected according to the LRT (p-value=0.05), using the data simulated under H0. The power prediction was defined as the fraction of the *N* data for which the LRT rejected H0 (p-value=0.05), using the data simulated under H1. The 95% confidence interval (CI) of both the type I error rate and the power predictions were computed assuming a binomial distribution.

The type I error and power predictions were computed using different cut-off values for the LRT: the nominal value issued from the *χ*^2^ distribution and the actual value computed with randomization test in order to calibrate the cut-off value to the actual data[19]. For each of the *N* datasets simulated under H1 a cut-off value needed to be computed. It was done via a randomization test of the *TRT* variable, breaking any existing correlation between the response and the treatment allocation. 1000 permuted datasets were fitted by both H0 and H1, resulting in a distribution of 1000 Δ*OFV* from which the 5th percentile was taken as the actual cut-off value (p-value=0.05). To limit the computational costs the randomization test was performed on 50 datasets instead of all 500. It resulted in 50 actual cut-off values for each combination of nested models and each approach, from which the mean, median and the mode were used as actual cut-off value for the prediction of type I error and power over the 500 datasets.

The accuracy of the drug effect estimate was computed using the final estimates of the H1 models fitted to the *N* data simulated with drug effect. For each model and each dataset, the estimated median drug effect 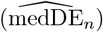 was computed according to Eqn. 11 (for two sub-populations and 1:1 randomization),

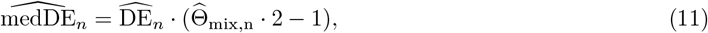

where 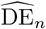 is the typical drug effect estimates in the dataset *n*, 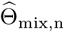 the mixture proportion estimate (set to 1 for STD). For the time independent drug models (*M*1 to *M*10 and *M*12), 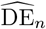 was equal to the typical value of the estimate, while for the time-dependent models 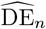 was evaluated at the last observation time (*t* = 3), using the typical estimates. The median drug effect estimates were compared to the true drug effect value used in the simulations (TDE = 9, with Θ_mix_ = 1 and DE = 9) to assess the accuracy of the approaches. The accuracy in the drug effect estimates was quantified with the relative estimation errors for each of the *N* simulated dataset with drug effect (*REE*_*n*_) as described in Eqn. 12:

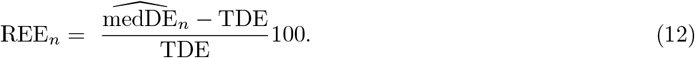

## 4 Results

After simulations, the median CFB between the two arms (ΔΔ*Response*) was 0 for the data simulated under H0. ΔΔ*Response* was randomly positive or negative across the *N* datasets. For the data simulated under H1, the median ΔΔ*Response* was equal to 10.2, representing 10% of the typical value of baseline. The range of ΔΔ*Response* difference in the data simulated with drug effect was [ − 4.65, 27.2], meaning that the response in the treated arm was lower in median than in the placebo arm for 3.3% of the 500 simulated dataset.

The type I error predictions using the nominal cut-off values for both NLMEM-based approaches, for the POC design (30 patients per arm) and the BIG design (120 patients per arm) are presented in Figure 1. On the thirteen models fitted to the datasets simulated without drug effect, the type I error rates were controlled for eleven models with IMA in both the POC and BIG designs, while they were controlled for three and two models with STD in POC and BIG designs respectively, as the 95% CIs covered the pre-defined value of 5%.

**Figure 1:**
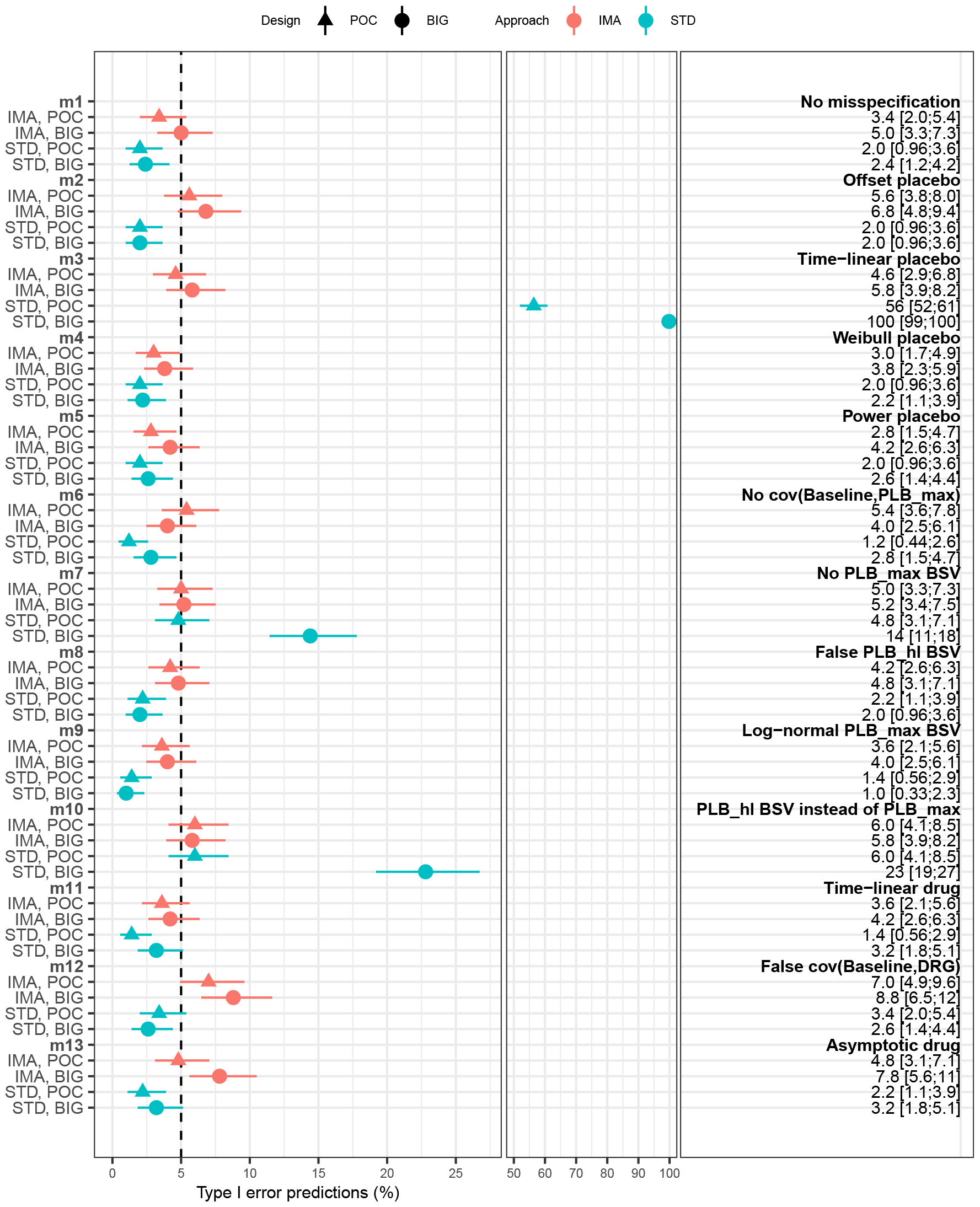
Type I error predictions using nominal cut-off values for IMA and STD, under the POC and BIG designs for each of the the thirteen models used for estimation. The error bars represents the 95% CI of the predictions based on 500 simulated dataset. The corresponding numbers are given on the right. The dashed vertical line represents the expected 5% type I error.

Using the non-misspecified model (*M*1), the type I error was controlled with IMA in both POC and BIG designs with a rate predicted at 5% for the latter. On the contrary, the predicted type I error rates were significantly lower than the expected 5% with STD in both designs. Regarding the performance of IMA with the misspecified models, the type I error rate was lower than expected for IMA for the models *M*4 and *M*5 in the POC design, while it was controlled in the BIG design. *M*4 and *M*5 were misspecified in the structure of the placebo model with respectively a Weibull and Power model. IMA had inflated type I error predictions for the models *M*12 and *M*13, overparameterized drug models in the BSV or the structural model respectively, in the BIG design. The type I error was controlled for both in the POC design even though higher than expected for *M*12.

Regarding STD, the overall poor performance in type I error control in the POC design were confirmed in the BIG design, showing that the results were independent to the sample size. Most of the type I error predictions with STD were significantly lower than the expected 5%. However highly inflated type I error rates were also observed for models *M*3 in both designs, and models *M*7 and *M*10 with the BIG design. *M*3 assumed a simpler time-linear placebo model, resulting in type I error predictions of 56% (POC design) and 100% (BIG design). *M*7 corresponded to the suppression of the PLB_max_ BSV, while *M*10 had this BSV suppression replaced by one in the PLB_hl_ parameter.

Figure 2 presented the type I error rates computed for the POC design using the nominal or the actual corrected cut-off value. The corrected value was obtained through 50 randomization tests (1000 randomizations each) on datasets simulated with drug effect. As some variability was observed across the 50 cut-off values, the mean, the median and the mode were used successively to calibrate the LRT (see Figure 5 in supplementary material). The three types of calibrations performed well with STD as the 95% CI of the type I error predictions covered the pre-defined value of 5% with most models. The best performances were observed using the median of the actual cut-off values, with a type I error control in all the models but *M*3, and predicted rates closer to the expected 5% (Figure 2). The LRT calibration decreased the type I error inflation for STD but did not result in a proper type I error control.

**Figure 2:**
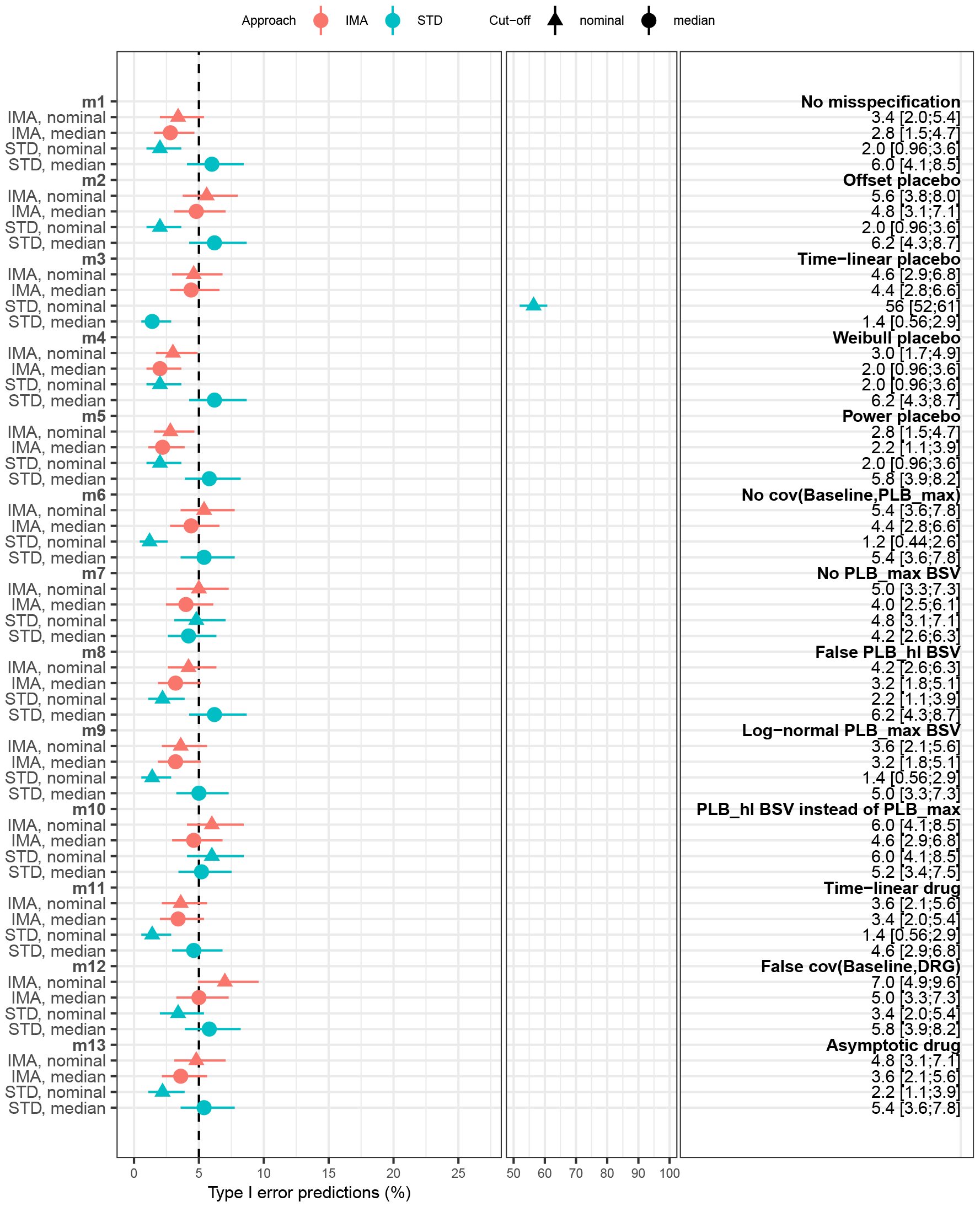
Type I error predictions for IMA and STD, under the POC design, at nominal or actual median (after LRT calibration) cut-off values for each of the the thirteen models used for estimation. The error bars represents the 95% CI of the predictions based on 500 simulated datasets. The corresponding numbers are given on the right. The dashed vertical line represents the expected 5% type I error.

Regarding IMA, the three types of LRT calibrations had very similar performance, all resulting in a decrease in the type I error predictions for all the models, even when the nominal values were lower than the expected 5% (see Figure 5 in supplementary material). Using the median of the actual cut-off values, the type I error rates of IMA were controlled in 10 models (Figure 2). The LRT calibration resulted in a controlled type I error rate for the model *M*12, i.e., only when the rate was higher than the expected 5% with the nominal cut-off. The LRT calibrations did not improve the type I error control when the predicted rates using the nominal cut-off were lower than the expected 5% (*M*4 and *M*5). With the calibrated LRT, the type I error dropped below controlled for the non-misspecified model (*M*1). Overall, with the exception of *M*3, the predicted type I error rates were slightly higher with STD than with IMA (Figure 2).

The power predictions under the POC design using the nominal and the median-calibrated cut-off values for STD and IMA with the thirteen estimation models are presented in Figure 3. The same results including the three types of LRT calibrations are presented in the supplementary material in Figure 5. The median of the power predictions ranged between [30-76]% for IMA and [28-100]% for STD using the nominal cut-off values. The usage of the median-calibrated cut-off values resulted in a small systematic decrease of the power prediction for IMA ([27-73]%) but had a bigger impact on the power rates for STD ([48-87]%).

**Figure 3:**
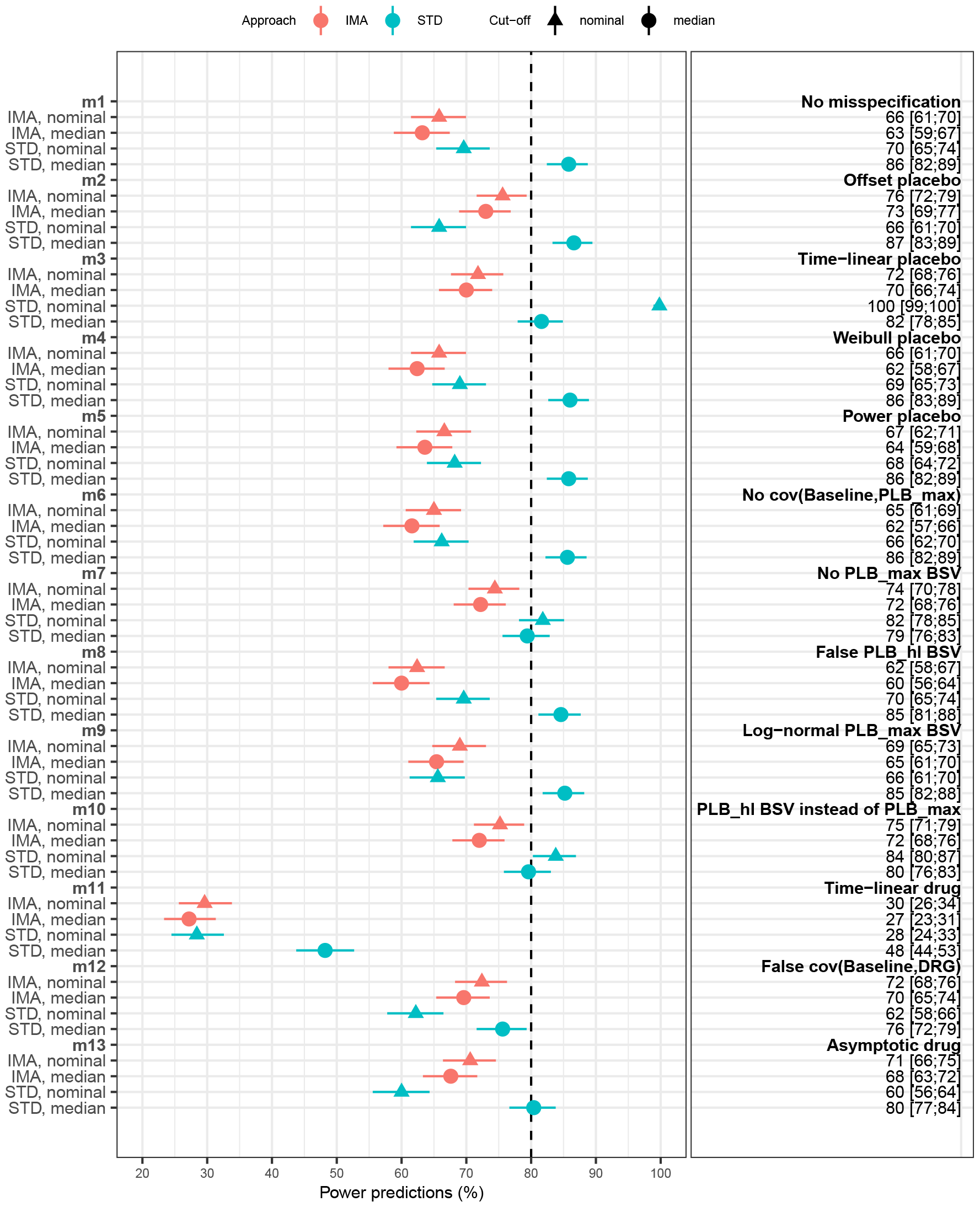
Power predictions for IMA and STD, under the POC design, at nominal or actual median (after LRT calibration) cut-off values for each of the the thirteen models used for estimation. The error bars represents the 95% CI of the predictions based on 500 simulated datasets. The corresponding numbers are given on the right. The dashed vertical line represents a visual reference at 80% power.

Using the nominal cut-off values, IMA and STD had similar performance for seven models, with overlapping 95% CI. Within the six remaining models, IMA had better power for *M*2, *M*12 and *M*13, while STD had better power for *M*3, *M*7 and *M*10. The LRT calibration resulted in a large improvement for STD for which the power after calibration was then systematically higher than the IMA predictions, even when based on the nominal cut-off. Comparing to the nominal cut-off based IMA power predictions, the median-calibrated STD power predictions were higher of 4 to 23 points across models. In *M*11 that assumed a time-linear drug model, the power predictions were very low both IMA and STD before and after LRT calibration.

Regarding the classical statistical analysis, the type I error rates were controlled for both, with predictions at 5.2 and 5.0% for the t-test and MMRM respectively (Table 3). The power predictions were 54% (95% CI [50;59]) for the t-test and 69% (95% CI [64;73]) for MMRM. The power of the t-test was significantly lower than the NLMEM-based approaches using nominal or median-calibrated cut-off values. For MMRM the power predictions were similar to the NLMEM-based approaches regardless of the calibration, with 95% CI overlapping. The power in STD after LRT calibration was significantly higher than all the other approaches (Table 3).

**Table 3:** Comparison of predictions (%) [95% CI] of type I error and power of LRT for IMA at nominal cut-off and after LRT calibration, STD at nominal cut-off and after LRT calibration, t-test and LRT of MMRM for the simulated POC design. Predictions and CI are based on 500 simulated datasets. Predictions for IMA and STD approaches are based on results with model *M*1 (same as simulations).

| Approach | Type I error (prediction % [95% CI]) | Power (prediction % [95% CI]) |
| --- | --- | --- |
| IMA | 3.4 [2.0;5.4] | 66 [61;70] |
| IMA + LRT calibration | 2.8 [1.5;4.7] | 63 [59;67] |
| STD | 2.0 [0.96;3.6] | 70 [65;74] |
| STD + LRT calibration | 6.0 [4.1;8.5] | 86 [82;89] |
| t-test | 5.2 [3.4;7.5] | 54 [50;59] |
| MMRM | 5.0 [3.3;7.3] | 69 [64;73] |

The accuracy in drug effect estimates was measured for the H1 of both IMA and STD for each of the 500 datasets simulated with drug effect under a POC design. The results are presented in Figure 4. The REE_*n*_ were similar between IMA and STD in most models, their median accross the *N* datasets were closer to 0 in IMA compared to STD. Differences were observed in the REE_*n*_ distribution for the model *M*3 for which IMA was less biased, and for *M*11 where STD was less biased. The highly positive STD REE_*n*_ values for *M*3 showed that estimates of the median drug effect were higher than expected and explained the related high predictions for both type I error and power. On the contrary the negative REE_*n*_ values in both IMA and STD explained the low power predictions of the model *M*11, as the median drug effect was underestimated.

**Figure 4:**
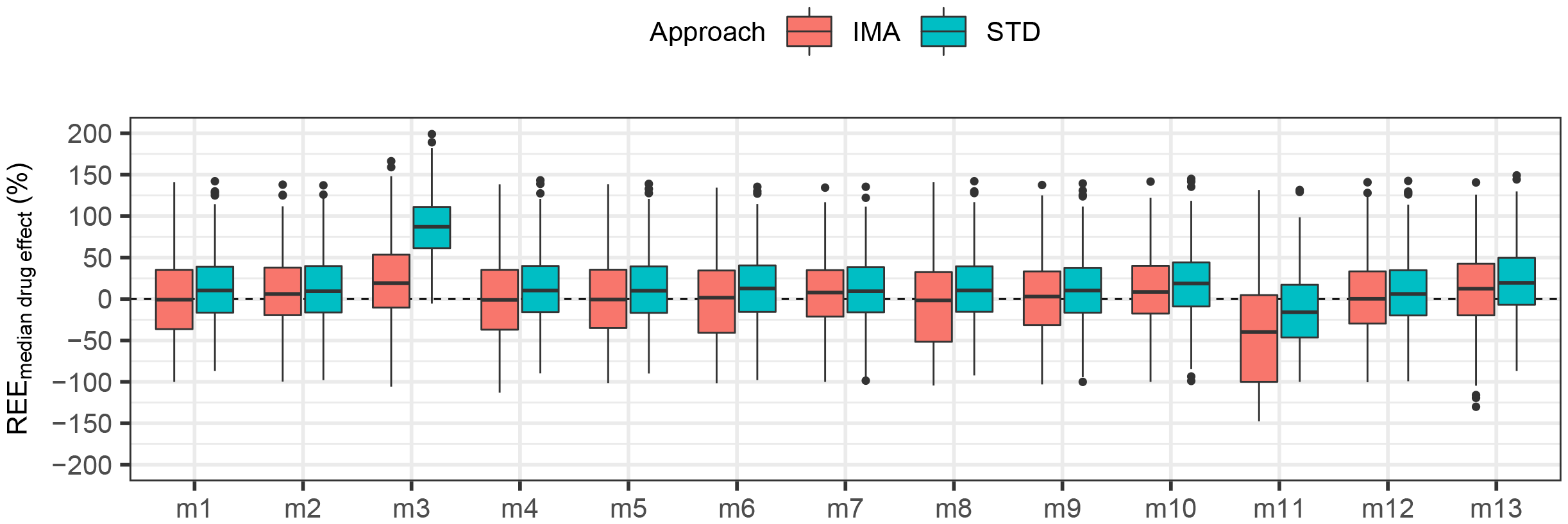
Accuracy of the drug effect estimate measured by the relative estimation errors (*REE*_*k*_) for each of the thirteen fitted models, under the POC design, in IMA and STD (n=500).

## 5 Discussion

IMA is a NLMEM-based approach using mixture models that previously showed promising performances to test for drug effect. IMA had good type I error control and unbiased drug effect estimates compared to STD in the context of balanced two-arms studies using real placebo data[12]. As NLMEM-based approaches already demonstrated interesting properties to increase the efficiency of the drug development process[1–7], this work aimed at extending the assessment of IMA and STD further, in cases where the extent of the model misspecification is known through a simulation study. A response model with or without drug effect was used as motivating example to allow the assessment of type I error, power, and bias in the drug effect estimates under various types of model misspecification. Their performances were also compared with more classical statistical approaches: t-test and MMRM.

The type I error when testing for drug effect was first assessed for IMA and STD, using the LRT with nominal cut-off values from the *χ*^2^ distribution (p-value=0.05), under two scenarios differing by the sample size. The POC design mimicked a traditional POC study with about 30 individuals per arm while the BIG design had a four times larger sample size to ensure asymptotic conditions for the LRT. For IMA in all thirteen models tried the two designs had type I error rates with overlapping CI, suggesting no impact of the sample size effect on the results. Similar results were observed for STD with four exceptions. The BIG design was associated with inflated and significantly higher type I error in all but one case: *M*3. In *M*3 the misspecification consisted in a reduced structural placebo model and the type I error was already inflated to a lower extend with the POC design, while in the two other cases (*M*7 and *M*10) the type I error was controlled with the POC design. Overall the IMA approach performed well with controlled type I error rates at the expected 5% value in all but two of the tested models. The type I error inflation were observed with the BIG design and involved the two models overparameterized in the drug component, either in the structural or the BSV model with addition of a covariance term. In a different context, using a pharmacokinetic (PK) model, with a different estimation method (FOCE with interaction) and heteroscedastic residual error models, the addition of a false covariance term also led to a systematic increase in type I error[6]. STD had poorly controlled type I error with predictions often below the expected 5% for most the tested models regardless of the design. Four exceptions showed type I error inflation instead: the model *M*3 with the two designs, for which the placebo model was reduced to a time-linear model, and the two models for which the BSV was omitted or simplified for the placebo model: *M*7 and *M*10. This inflation could be explained by a loss of flexibility to capture the observed data through the placebo model, compensated by the drug model. It is however interesting to observe the inflation under the theoretically better asymptotic conditions.

A calibration of the LRT via a randomization test (1000 samples) was performed. Randomization tests are a recommended practice to decide on the statistical significance of a covariate in NLMEM with real data, especially in cases where the asymptotic conditions are not met (e.g small sample sizes)[6]. Ideally such a test should be carried out for each of the thirteen models and each of the 500 datasets. To reduce the computational costs, the randomization test was performed on a restricted number of 50 datasets for each model. Hence instead of having one calibrated cut-off value per dataset, different statistics of the 50 cut-off values were computed and used (mean, median and mode). The calibration had no significant impact on IMA for which the CI around the predicted type I error rates were systematically overlapping. However the need for calibration was limited in IMA as the type I error was already well controlled in most models with the nominal cut-off values. On the contrary for STD the CI did not overlap for 7 out of 13 models, suggesting a stronger impact of the LRT calibration. A direction was observed also: the calibrated test resulted in higher type I error rates, bringing most of the models to controlled rates. With the exception of model *M*3 for which it led to a drastic type I error reduction, from 56% to below controlled. The randomization test performed well in STD as type I error was controlled afterwards in most models.

The data simulated with drug effect allowed to assess the approaches both in terms of power and bias in the drug effect estimates under the POC design. The calibration had no significant impact on IMA for which the CI around the predicted power rates were systematically overlapping. These results are coherent with what was observed for the type I error rates. On the contrary, for STD the CI of the power rate prediction overlapped only for 3 models, suggesting a strong calibration impact, improving the power for all the other models but one (*M*3). These results were also coherent with the overall type I error increase observed after the LRT calibration. The median of the drug effect estimates was similar for IMA and STD, consistently slightly closer to 0 for IMA. Both IMA and STD were biased similarly with the misspecified structural drug model (*M*11), but IMA was less biased in the case of an underparameterized placebo model (*M*3). These poor bias performances are associated with atypical power behavior. The power for STD with *M*3 was of 100% (highest type I error of 56%). *M*11 on the contrary had the lowest power for both approaches, regardless of calibration. Despite these similar performances in the estimates of the drug effect size, differences in power predictions were observed. Using nominal cut-off values for the LRT, power predictions with IMA were higher or similar to STD in most of the models. When using median-calibrated cut-off for the LRT, STD had systematically better power performances than the best IMA performances.

Type I error predictions were slightly higher in STD after LRT calibration than in IMA but this difference could only explain a small part of the differences observed in terms of predicted power. In comparison, both the t-test and the MMRM approaches had controlled type I error rates. The power observed with the t-test was lower than the two NLMEM-based approaches power predictions, confirming previous results[1]. However, the usage of the more sophisticated MMRM analysis, based on linear mixed-effects model, led to power predictions of the same magnitude as IMA, hence significantly lower than what was observed for STD after LRT calibration.

Wählby et al. previously assessed the type I error of the LRT in pharmacometric model-based analysis to test for the inclusion of an extra unnecessary parameter such as a covariate, a random effect or a covariance with a PK model describing simulated continuous data[6, 19] or simulated categorical data[5]. The addition of a false covariate in a PK model using the FOCE estimation method led to a type I error higher than expected when residual error model was heteroscedastic (exponential or proportional), while only a slight increase of type I error was observed with a homoscedastic (additive) error model[19]. The addition of a false drug effect (fixed effect only) in the logistic model using the LAPLACE estimation resulted in a type I error close to the expected nominal value[5]. The type I error was predicted lower than expected when a false random effect for BSV of the drug effect was tested in the logistic model[5]. In the present work fitting the non misspecified placebo model with a drug effect component on data simulated without drug effect resulted in expected controlled type I error rate in IMA, while it was significantly lower in STD. In this latter approach LRT was performed for a false fixed effect parameter and a false random effect describing the drug effect. Overall, the test of a false random effect for BSV of the drug effect could explain the systematically lowest predictions of type I error in STD, as shown previously for another type of model (logistic) and estimation method (LAPLACE)[5].

Aoki et al. compared model-averaging and model selection in a simulation study in the context of dose finding trials[11]. They showed both methods achieved similar probabilities to find the correct dose and a similar type I error rate that was slightly lower than the expected value when using data simulated without drug effect. No formal hypothesis test was performed as a type I error was defined as the selection of one of the tested dose by the method. Using a sampling case bootstrap procedure on the initial dataset to take into account the drug-response model uncertainty, both methods achieved a higher probability to find the correct dose. Buatois et al. also compared model-averaging to model selection in simulated dose finding trials. They showed that model-averaging resulted in better predictive performances with more accurate predictions of the response and better prediction of the minimal effective dose[20]. No type I error or power rates were reported in the work as the selection of the drug model was not based on hypothesis testing. In both works model-averaging was performed using a pre-defined set of drug effect models, assuming a well specified placebo model. The use of model-averaging in a context of hypothesis testing in NLMEM has not yet been assessed. The present work assessed how misspecifications of the placebo effect model could impact the type I error and power predictions. It highlighted for example the sensibility of STD to such misspecifications, resulting in strongly inflated type I error rates. A model-averaging approach using different set of placebo and drug effect models could achieve better predictive performances and would be highly suitable to pre-specify pharmacometric model-based analysis.

IMA confirmed its promising results, especially in terms of type I error control. Its performances could be assessed further, to unbalanced designs, other types of data (categorical data or count data), as well as different residual models, or more complex cases of model misspecification. It could also be interesting to expand the work to a larger set of mixtures to have an approach that would allow model pre-specification for many analysis. Note that in the case of a larger number of mixtures, the functions used in IMA in the present analysis should be revised and could require heavy mathematical work to properly implement the mixture models, for example at least to constrain all the probabilities. Previous work combining bootstrap methods to traditional model-averaging resulted in better performances than model-averaging only[11]. It could be interesting to assess how bootstrap methods could be helpful in the IMA approach.

## 6 Conclusion

The present work assessed the performances of IMA and STD when testing for drug effect in the context of two-arms balanced studies with simulated data. Their performances were compared to traditional statistical approaches: t-test or MMRM. IMA confirmed its encouraging performances (type I error control and unbiased drug estimates) observed previously on real placebo data. Using simulated data with and without drug effect IMA had good type I error control, showed good power properties and unbiased drug estimates. However the power performances were outperformed by the standard approach when calibrated properly, likely due to the additional parameter estimate required for IMA. IMA is a promising NLMEM-base approach for hypothesis testing of the drug effect and could be used in the future, after further evaluations, as primary analysis in confirmatory trials.

## 8 Supplemental material

**Figure 5:**
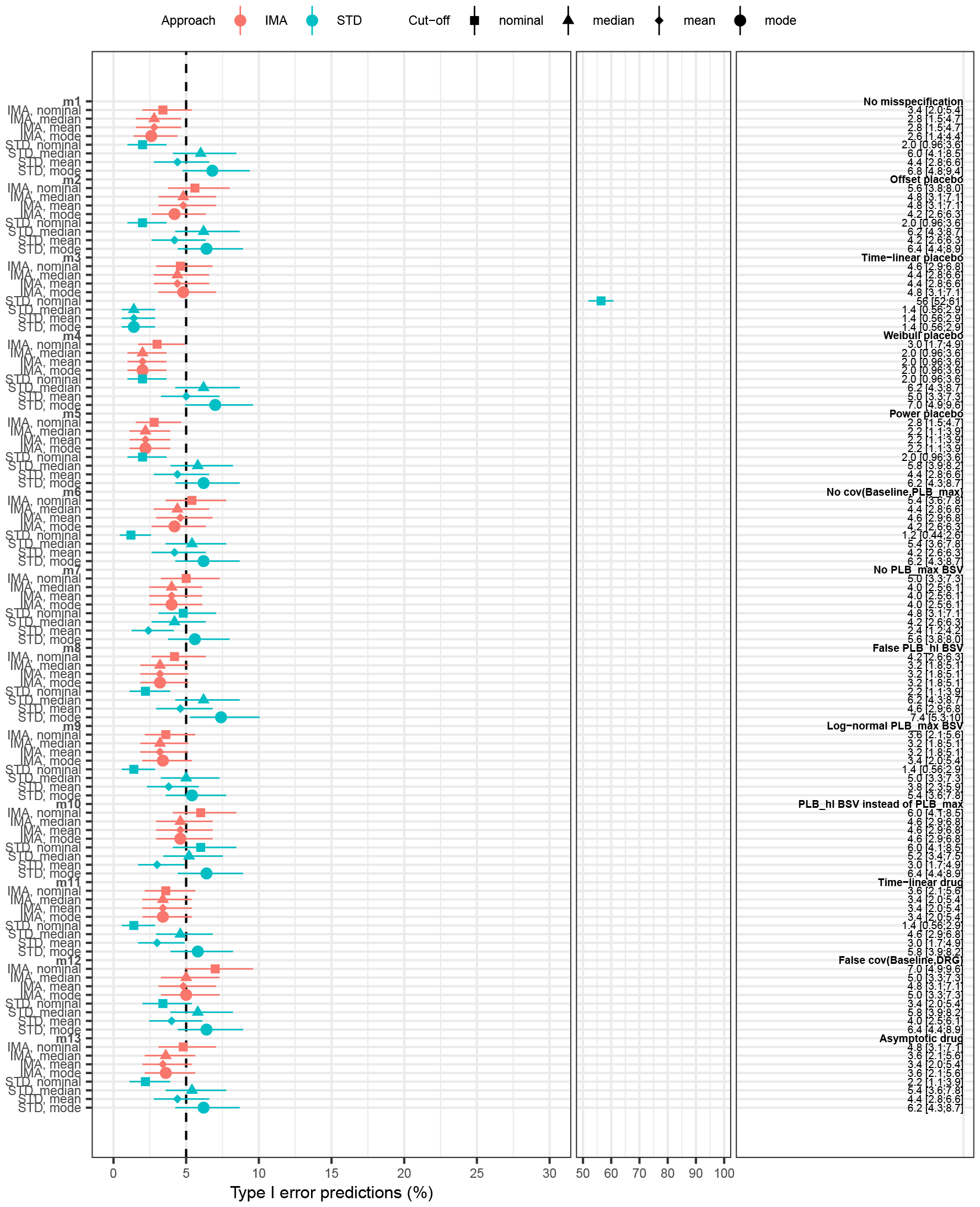
Type I error predictions for IMA and STD under the POC design, using nominal or actual (after LRT calibration) mean, median or mod cut-off values for each of the the thirteen models used for estimation. The error bars represents the 95% CI of the predictions based on 500 simulated datasets. The corresponding numbers are given on the right. The dashed vertical line represents the expected 5% type I error.

**Figure 6:**
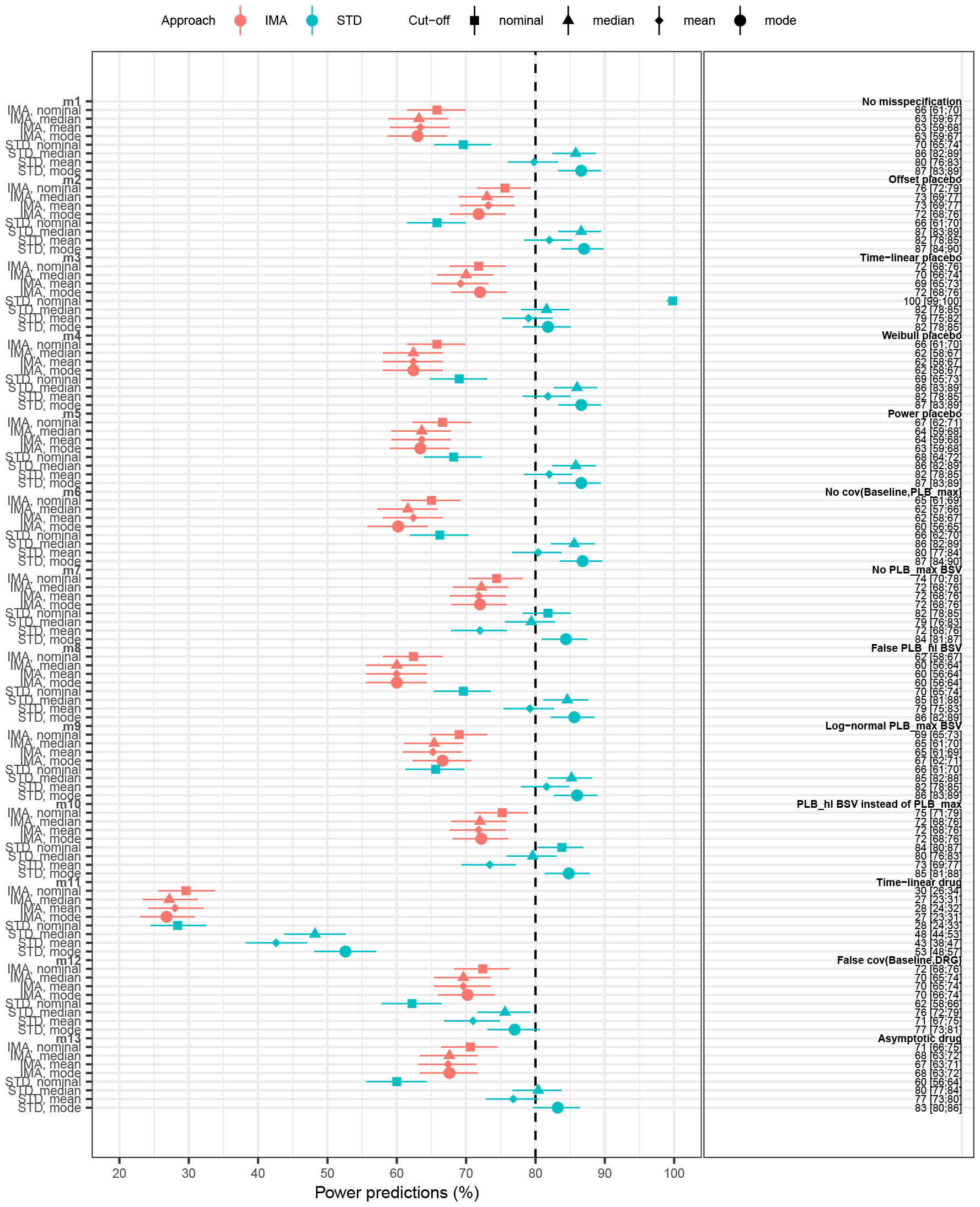
Power predictions for IMA and STD under the POC design, using nominal or actual (after LRT calibration) mean, median or mod cut-off values for each of the the thirteen models used for estimation. The error bars represents the 95% CI of the predictions based on 500 simulated dataset. The corresponding numbers are given on the right. The dashed vertical line represents a visual reference at 80% power.

**Figure 7:**
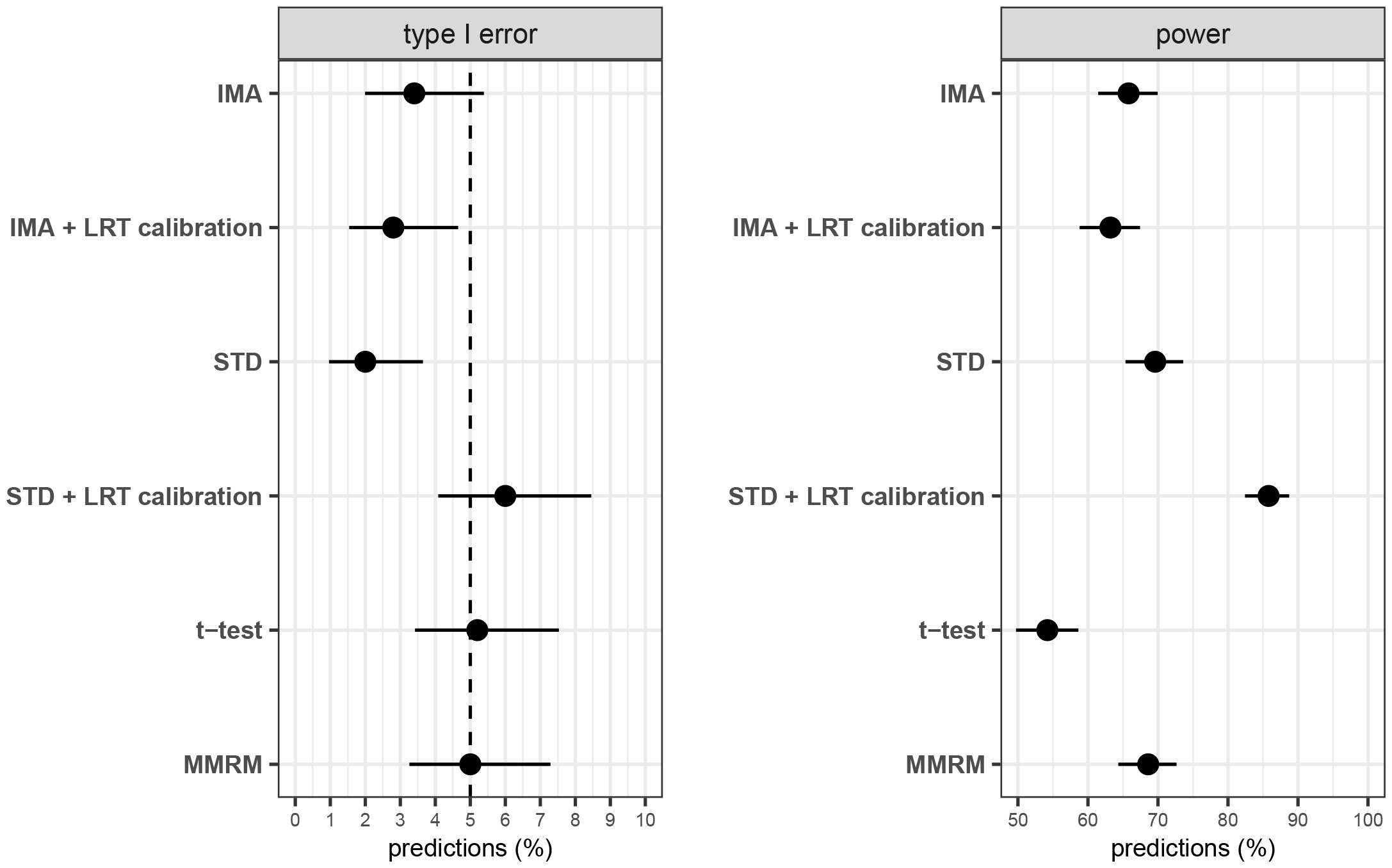
Comparison of type I error (left) and power predictions (right) for LRT in IMA at nominal cut-off values (IMA) and after LRT calibration (IMA + LRT calibration), STD at nominal cut-off values (STD) and after LRT calibration (STD + LRT calibration), t-test and LRT in MMRM analysis (MMRM). Error bars represents the 95% CI of predictions based on 500 simulated dataset. Dashed vertical line represents the expected 5% type I error.

